# Cyrene: A Novel Geroprotective Compound that Extends Lifespan and Healthspan in *C. elegans* and *Drosophila*

**DOI:** 10.1101/2025.08.01.668202

**Authors:** Abdelrahman AlOkda, Shweta Yadav, Alain Pacis, Andrey A. Parkhitko, Jeremy M. Van Raamsdonk

## Abstract

As aging is the primary risk factor for many chronic diseases, geroscience aims to target aging to delay age-related decline. Here, we identify Cyrene (dihydrolevoglucosenone), a sustainable, biocompatible solvent, as a novel geroprotective compound. Cyrene extends lifespan and healthspan in *C. elegans*, improving locomotor function and resistance to oxidative, thermal, osmotic, genotoxic, and proteotoxic stress. It also confers protection in neurodegenerative models of Alzheimer’s, Parkinson’s, and Huntington’s disease. Cyrene is effective when delivered during development or early adulthood and requires administration before day 8 to extend longevity. Its benefits are independent of bacterial metabolism and partially independent of the FOXO transcription factor DAF-16. Importantly, Cyrene also extends lifespan and enhances oxidative stress resistance in *Drosophila melanogaster*, demonstrating cross-species efficacy. These findings identify Cyrene as a novel geroprotective compound that promotes longevity, resilience, and neuroprotection. Conservation across species supports future work to dissect molecular mechanisms and test its potential in mammals.

## Introduction

Aging is characterized by progressive deterioration in stress tolerance, proteostasis, and tissue integrity leading to increased susceptibility to neurodegeneration, metabolic dysfunction, and mortality. Aging is the primary risk factor for chronic diseases and functional decline across species ^1^. Rather than developing treatments for each chronic disease individually, geroscience aims to develop treatments that target the aging process, with the goal of delaying or preventing multiple chronic diseases simultaneously. A central goal in geroscience is to identify interventions that not only extend lifespan but also preserve physiological function with minimal trade-offs. Several small molecules, including metformin, rapamycin, and resveratrol, have been shown to extend longevity and protect against chronic disease in model organisms ^2,3^. These compounds act through nutrient-sensing, stress-response, or mitochondrial signaling networks.

In searching for novel geroprotective compounds, which extend lifespan and protect against chronic diseases, we found that Cyrene (dihydrolevoglucosenone) extends lifespan in wild-type worms, prompting a systematic investigation into its biological activity. Cyrene is a polar, water-miscible solvent derived from cellulose and was originally developed as a sustainable alternative to toxic solvents such as N,N-dimethylformamide (DMF) and N-methyl-2-pyrrolidone (NMP) ^4,5^. While extensively characterized for chemical and industrial applications, Cyrene has not been studied as a bioactive compound. It was presumed to be inert in biological systems, and to our knowledge, no studies prior to this work had evaluated its effects on aging, stress response, or organismal function.

In this study, we characterize the phenotypic effects of Cyrene in two genetic model organisms: *C. elegans* and *Drosophila melanogaster*. We show that exposure to Cyrene extends lifespan without major developmental or reproductive trade-offs, preserves neuromuscular function in aging animals, enhances resistance to diverse stressors, and mitigates functional decline in models of neurodegeneration. The beneficial effects of Cyrene are dose-dependent, time-sensitive, independent of bacterial metabolism, and partially independent of DAF-16. Together, these findings demonstrate that Cyrene is a novel, geroprotective compound with lifespan-extending properties that are conserved across species.

## Results

### Cyrene extends lifespan and healthspan with minimal developmental or reproductive trade-offs

Research from our laboratory and others has shown that genes and compounds that extend lifespan can also protect against neurodegenerative diseases ^2,6,7^. In searching for novel geroprotective compounds, we found that Cyrene significantly extends lifespan in *C. elegans*. Having identified Cyrene as a promising candidate, we exposed worms to concentrations ranging from 0.1% to 4% (v/v) to define its optimal dose and exposure window for lifespan extension in *C. elegans*. Treatment with Cyrene was initiated at either the L1 stage to maximize exposure to the compound, or in young adults. We found that Cyrene increased lifespan at concentrations ranging from 0.1% to 2% with maximal benefit observed at 1% (**Figure 1A**). At concentrations of 2.5% or above, lifespan was decreased, reflecting potential toxicity at these doses. The ability of Cyrene to increase lifespan was consistent whether exposure began at the L1 larval stage or in early adulthood (**Figure 1A,B**).

**Figure 1.**
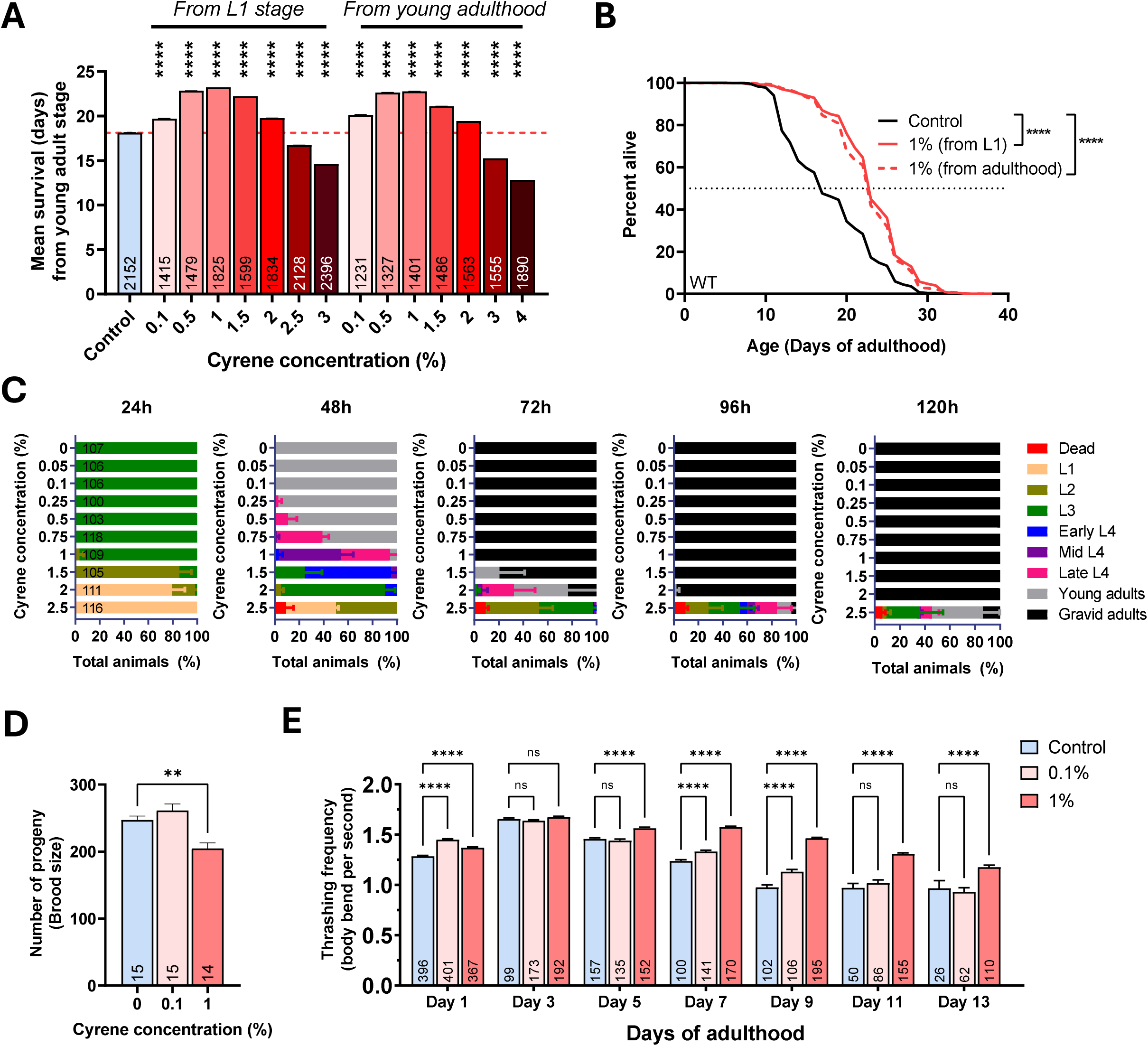
Cyrene extends lifespan and healthspan in *C. elegans* with minimal developmental or reproductive trade-offs. Cyrene treatment from either larval (L1) or young adult stages led to significant improvements in longevity and preserved function without markedly affecting growth or reproduction. (A) Survival curves show that 1% Cyrene extended lifespan when administered from either the L1 stage or young adulthood, with maximal effect observed in both conditions (B) Mean lifespan of worms treated with varying concentrations of Cyrene starting from either the L1 stage or young adulthood. The optimal concentration (1% v/v) significantly extended lifespan (*p* < 0.0001 vs. control). (C) Post-embryonic developmental timing (24–120 h) under Cyrene exposure; 1% treatment caused only a minor delay. Full statistical comparisons are provided in **Supplemental Table S1**. (D) Fecundity analysis (total progeny count) under 1% Cyrene treatment, showing minimal reproductive impact. (E) Age-associated decline in thrashing frequency was attenuated in Cyrene-treated animals, suggesting improved healthspan. Error bars represent standard error of the mean (SEM). Sample sizes (n-values) are indicated within each bar. Statistical analysis was performed using the log-rank test with Bonferroni’s multiple comparisons test for panel A, one-way ANOVA with Dunnett’s multiple comparisons test for panels B and D, and two-way ANOVA with Šidák’s multiple comparisons test for panel E. ns = not significant, ***p < 0.001, ****p < 0.0001 from control.

In order to further optimize our Cyrene treatment protocol, we compared two delivery methods: applying Cyrene on top of the agar after nematode growth medium (NGM) plates have been poured versus direct incorporation of Cyrene into the NGM agar prior to pouring. While both approaches extended lifespan, the on-top method produced a more robust and reproducible effect (**Figure S1**). The reduced efficacy of the in-agar method may reflect compound degradation, bacterial metabolism, or limited diffusion from the agar matrix into the worm’s microenvironment.

To determine the extent to which the increase in lifespan from Cyrene treatment is associated with developmental and reproductive trade-offs, we measured post-embryonic developmental time (from hatching to adulthood) and fertility (total brood size). Consistent with the results from the lifespan experiment, concentrations above 2% of Cyrene resulted in marked development delays (**Figure 1C**). At the optimum dose for lifespan of 1% Cyrene, worms developed with a slight (∼2–3 hours) delay (**Figure 1C**). At 0.1% Cyrene, the post-embryonic development time was indistinguishable from the 0% control while still extending longevity. In examining the effect of Cyrene on fertility, we found that there was no effect of treatment with 0.1% Cyrene and only a mild reduction in total progeny at 1% Cyrene (**Figure 1D**).

To evaluate the effect of Cyrene on healthspan, we assessed age-associated neuromuscular decline by measuring the rate of movement in liquid (thrashing rate) ^8^. While control animals showed the expected progressive decrease in movement with age, animals treated with 1% Cyrene retained significantly higher locomotor activity between days 5 and 13 of adulthood (**Figure 1E**). Treatment with 0.1% Cyrene also preserved the rate of movement in aging worms but to a lesser extent than 1% Cyrene (**Figure 1E)**. The preservation of motor function with age suggests that Cyrene not only increases lifespan but also delays the onset of physiological decline. Combined, these results indicate that Cyrene enhances both lifespan and healthspan without incurring major developmental or reproductive costs.

### Cyrene extends lifespan independently of bacterial metabolism

Several small molecules extend *C. elegans* lifespan not by acting on the worm, but by impairing bacterial growth, altering bacterial metabolism, being metabolized by bacteria or through changes in bacterial virulence ^9^. As OP50 bacteria are mildly pathogenic to *C. elegans* and worms have been shown to live longer when grown on dead bacteria or under dietary restriction ^10–13^, we assessed whether Cyrene has bactericidal activity at lifespan-extending concentrations. We conducted a dose-response assay measuring *E. coli* OP50 growth in the presence of Cyrene. While Cyrene decreased bacterial survival at concentrations above 2%, there was no effect on bacterial viability at lower concentrations including the optimal concentration for longevity of 1% Cyrene (**Figure S2**). This suggests that the observed longevity effects are not a consequence of bacterial growth inhibition.

To further assess whether Cyrene’s longevity effects depend on the microbial environment, we conducted lifespan assays using non-replicating bacterial food sources. Worms were fed either UV-irradiated *E. coli*, which are replication-deficient but retain some metabolic activity, or paraformaldehyde (PFA)-fixed *E. coli*, which are metabolically inactive and fully non-viable ^14^. In both conditions, Cyrene significantly extended worm lifespan (**Figure 2A–D**). These findings suggest that Cyrene promotes longevity through a direct effect on *C. elegans*, rather than via modulation of bacterial replication or metabolism.

**Figure 2.**
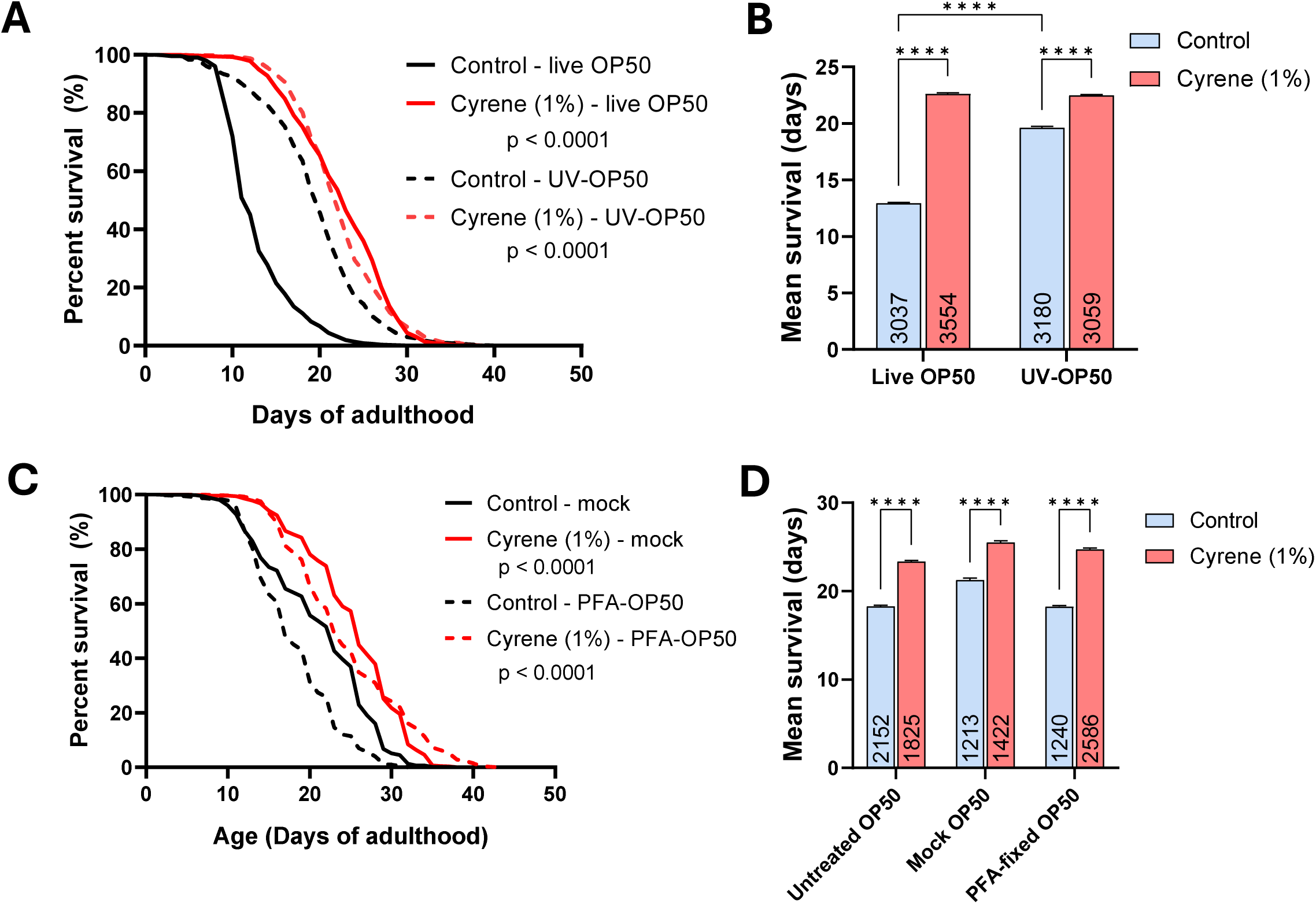
Cyrene extends lifespan independent of bacterial metabolism. Cyrene treatment significantly increased *C. elegans* lifespan across multiple bacterial conditions, suggesting that its pro-longevity effects are host-directed rather than microbiome-mediated. (A) Survival curves show that Cyrene extended lifespan when administered on either live or UV-irradiated *E. coli* OP50, the latter being replication-deficient but metabolically active. (B) Corresponding mean lifespan values confirm that Cyrene’s effect is independent of bacterial viability. (C) Lifespan extension was similarly observed when worms were fed Cyrene-treated OP50 grown in LB, mock-treated OP50 resuspended in PBS, or paraformaldehyde-fixed (metabolically inactive) OP50. (D) Corresponding mean lifespan values show no significant attenuation of effect across these bacterial states. These findings support a direct effect of Cyrene on the worm, independent of microbial metabolism or proliferation. Error bars indicate SEM. Sample sizes (n-values) are indicated within each bar. Statistical analyses were performed using a log-rank test with Bonferroni’s multiple comparisons test in panels A and C, and a two-way ANOVA with Tukey’s multiple comparisons test in panel B and D. ****p<0.0001.

### Cyrene enhances biological resilience throughout the life course

We and others have shown that stress resistance is positively correlated with aging ^15^ at least partially due to same genetic pathways contributing to both phenotypes ^16^. However, in some cases these phenotypes can be experimentally dissociated ^17–19^. To determine whether Cyrene promotes resilience, we evaluated its effect across a range of exogenous stressors. Worms were treated with Cyrene from the L1 stage and challenged with environmental or chemical insults at different adult timepoints. For most assays, we chose to examine worms at day 1, 3, 5, 7, 9 and 11 of adulthood, as Cyrene-treated and control worms start dying at different rates beyond day 11. Day 1 adults refer to young adults 2–3 hours post L4–adult molt. Importantly, Cyrene was withdrawn during and after stress exposure, allowing us to assess sustained, treatment-induced adaptations and avoid any direct interactions between Cyrene and the chemicals or bacteria involved in inducing the stress.

Worms treated with Cyrene maintained the ability to survive sub-lethal heat stress (35°C for 3 hours followed by 21 hours of recovery) in mid-to-late adulthood while heat stress resistance in control worms progressively declined with age (**Figure 3A**). Thermotolerance of the Cyrene-treated animals at day 6-11 of adulthood was equivalent to Cyrene-treated animals at day 3 of adulthood and significantly greater than control worms. We observed a similar preservation of heat stress resistance in Cyrene-treated worms under harsher heat stress conditions (35°C for 6 hours followed by 18 hours of recovery; **Figure S3A**).

**Figure 3.**
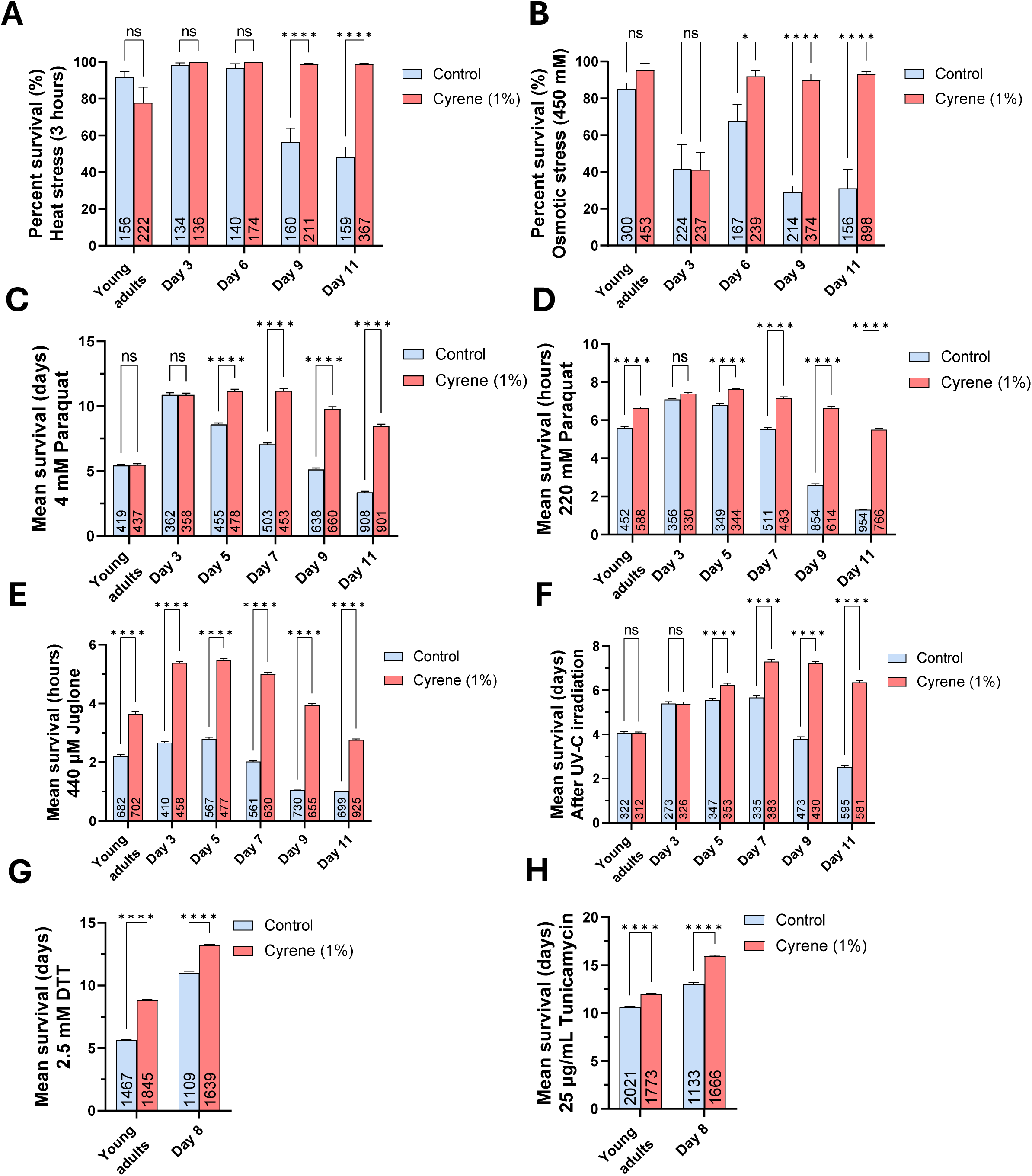
Cyrene enhances stress resilience in *C. elegans* across multiple acute and chronic stress paradigms. Cyrene-treated animals exhibited improved survival under diverse stress conditions. This included: (**A**) Age-dependent protection against heat stress (35°C for 6 h followed with 18 h recovery period), (**B**) hypertonic stress induced 500 mM NaCl (48 h), (**C**) chronic oxidative stress induced by 4 mM paraquat, (**D**) acute oxidative stress induced by 220 mM paraquat and **(E**) 440 μM juglone. Cyrene also increased resistance to (**F**) genotoxic stress induced by UV-C irradiation (0.1 J/cm²), (**G**) thiol-reductive stress by 2.5 mM Dithiothreitol (DTT), and (**H**) tunicamycin-induced ER-stress (25 µg/mL). These findings suggest that Cyrene broadly promotes cellular stress resilience through multiple mechanisms. Error bars represent the standard error of the mean (SEM). Sample sizes (n-values) are indicated within each bar. Statistical analyses were performed using two-way ANOVA with Šidák’s multiple comparisons test. ns = not significant, **p < 0.01, ****p < 0.0001.

As in the heat stress assay, Cyrene treated worms were able to maintain their peak resistance to osmotic stress from day 6 to 11 of adulthood, while control worms exhibited a progressive decline (450 mM NaCl; **Figure 3B**). Under osmotic stress, Cyrene-treated worms survived significantly longer than control worms at days 1, 6, 9 and 11 of adulthood (450 mM NaCl; **Figure 3B**). We observed a similar protective effect of Cyrene under harsher osmotic stress conditions (500 mM NaCl; **Figure S3B**).

Next, we examined the ability of Cyrene to protect against oxidative stress through exposure to either paraquat (chronic 4 and 10 mM, acute 220 mM) or juglone (acute 140, 260 and 440 µM). Cyrene conferred protection against both chronic (4 mM paraquat, 10 mM paraquat; **Figures 3C**, **S3C-D**) and acute paraquat exposure (220 mM paraquat; **Figures 3D**, **S4A**). This protection emerged from day 5 of adulthood onward and was strongest in late adulthood. Similarly, Cyrene-treated worms exhibited robust protection against juglone-induced oxidative stress across all ages (140, 260 and 440 µM juglone; **Figures 3E**, **S4B**).

We also found that Cyrene improved survival following UV-C irradiation, a model of genotoxic stress. The protective effect of Cyrene was age-dependent, becoming apparent after day 5 of adulthood and most pronounced by day 11 (0.1 J/cm²; **Figure 3F**, **S5A**). Finally, Cyrene improved resistance to both thiol-induced reductive stress (2.5 mM dithiothreitol; **Figures 3G, S5B-C**) and tunicamycin-induced endoplasmic reticulum (ER) stress (25 µg/mL tunicamycin; **Figures 3H, S5D-E**).

Together, these findings demonstrate that Cyrene treatment protects against multiple exogenous stressors indicating that it is not only a lifespan-extending compound but also extends healthspan. For all of the stresses we examined, Cyrene prevented or markedly reduced the aging-associated decline in resistance to stress ^20^ such that the maximum increase in stress survival was observed in the oldest worms.

### Cyrene preserves locomotor function in *C. elegans* models of neurodegenerative diseases

Our previous research and work from other laboratories have shown that genes and compounds that extend lifespan can also be neuroprotective in animal models of neurodegenerative diseases including Alzheimer’s disease (AD), Parkinson’s disease (PD), Huntington’s disease (HD), and Amytrophic Lateral Sclerosis (ALS) ^2,6,7^. To determine whether Cyrene treatment is beneficial in *C. elegans* models of age-associated neurodegenerative disease, we treated worm models of AD, PD and HD with Cyrene and quantified the ability of Cyrene to ameliorate deficits in movement at day 1, day 8 and day 12 of adulthood by measuring the rate of movement in liquid (thrashing rate). These strains express human disease-causing proteins including the main proteins implicated in AD (tau, Aβ₁₋₄₂), PD (α-synuclein, Leucine-rich repeat kinase 2 (LRRK2)), and HD (expanded polyglutamine protein) under neuronal, muscle or ubiquitous promoters, modeling key features of AD, PD and HD.

At day 1 of adulthood, we found that all of the untreated worm models of neurodegenerative diseases exhibited decreased movement compared to untreated wild-type worms (**Figure 4A**). While Cyrene treatment at this age resulted in a small decrease in movement in wild-type worms, it significantly increased movement in two of the three AD models, all three PD models and both HD models. At day 8 of adulthood, Cyrene significantly increased movement in all eight models of neurodegenerative disease tested (**Figure 4B**). Finally, at day 12 of adulthood, all strains showed decreased movement compared to younger ages. Nonetheless, treatment with Cyrene significantly improved movement in all three AD models, all three PD models and one of two HD models (**Figure 4C**).

**Figure 4.**
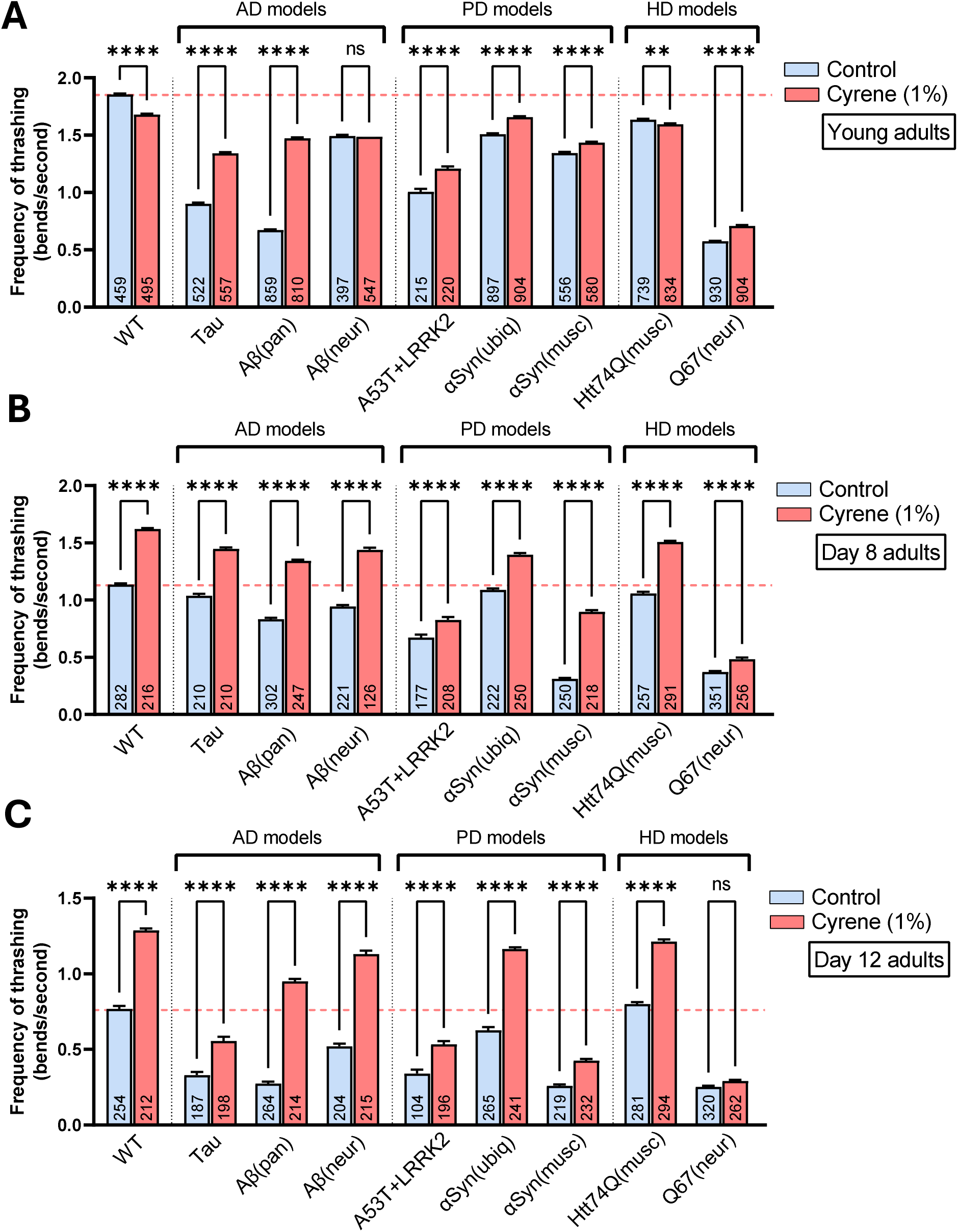
Cyrene rescues mobility deficits in *C. elegans* models of neurodegenerative diseases. Thrashing frequency was assessed at three timepoints: (**A**) young adulthood, (**B**) day 8, and (**C**) day 12 of adulthood. Strains included models of **Alzheimer’s disease** (AD)—<u>AD-Tau</u> CK10 (*aex-3p::tau[V337M]*), <u>AD-Aβ(pan)</u> CL2355 (*snb-1p::Aβ_1–42_*), and <u>AD-Aβ(neur)</u> GRU102 (*unc-119p::Aβ_1–42_*); **Parkinson’s disease** (PD)—<u>PD-A53T+LRRK2</u> JVR182 (*dat-1p::α-syn[A53T]; dat-1p::LRRK2[G2019S]*), <u>PD-αSyn(ubiq)</u> JVR389 (*eft-3p::α-syn[WT]*), and <u>PD-αSyn(musc)</u> NL5901 (*unc-54p::α-syn[WT]*); **and Huntington’s disease** (HD)—<u>HD-Htt74Q(musc)</u> MQ1698 (*unc-54p::Htt-74Q*) and <u>HD-Q67(neur)</u> AM716 (*rgef-1p::67Q*). Cyrene treatment increased thrashing frequency across all strains and timepoints, suggesting broad protective effects. Full genotype descriptions and disease associations are provided in the *Methods*. Error bars represent standard error of the mean (SEM). Sample sizes (n-values) are indicated within each bar. Statistical analyses were performed using one-way ANOVA with Šidák’s multiple

Importantly, Cyrene improved movement in neuron-specific models, muscle-restricted models and ubiquitous models, suggesting that its protective effects are not confined to a single tissue. In addition, the fact the Cyrene protects against the toxic effects of multiple different disease-causing proteins from multiple different diseases indicates that Cyrene is exerting a general beneficial effect on healthspan and may be able to protect against multiple diseases simultaneously.

### Cyrene treatment during development or adulthood extends lifespan

To define the temporal requirements for Cyrene’s pro-longevity effect, we systematically varied the timing and duration of treatment. Worms were exposed to Cyrene beginning at different stages from L1 to day 12 of adulthood and maintained on Cyrene plates until death (**Figure 5A**). We also examined the effect of Cyrene treatment beginning at L1 and stopping Cyrene treatment at different ages from young adulthood to day 12 of adulthood (**Figure 5A**).

**Figure 5.**
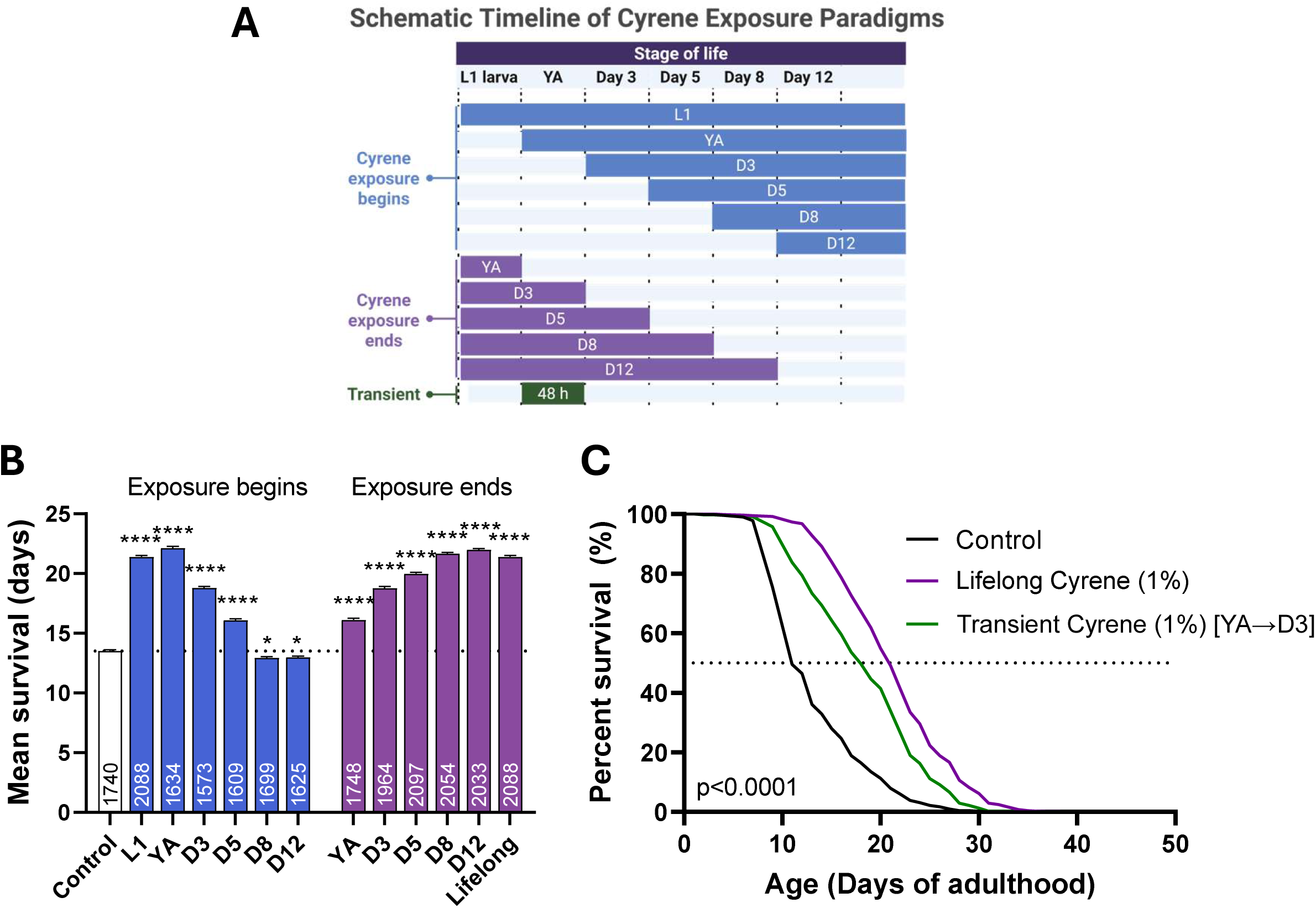
Early-life exposure to Cyrene is sufficient to extend lifespan. Lifespan assays were performed under temporally restricted exposure paradigms to determine the stage-specific requirements for Cyrene’s geroprotective effects. (A) In the “Cyrene exposure begins” paradigm, animals were treated with 1% Cyrene starting at defined life stages (L1, young adult [YA], Day 3, Day 5, Day 8, or Day 12) and maintained on treatment thereafter. In the “Cyrene exposure ends” paradigm, worms were exposed to Cyrene from L1 and transferred to untreated plates at the same respective timepoints. A separate group received only transient exposure (48 hours from YA to Day 3). (B) Mean lifespan data reveal that Cyrene’s longevity benefits were maximized when exposure began early and was maintained until at least Day 8 of adulthood, while treatments initiated at later stages produced progressively smaller effects. (C) Survival curves demonstrate that a brief, early-life exposure (YA to Day 3) was sufficient to induce significant and lasting lifespan extension, indicating a critical early-life window of action. Error bars represent the standard error of the mean (SEM). Sample sizes (n-values) are indicated within each bar. Statistical analyses were performed using one-way ANOVA with Dunnet’s multiple comparisons test for panel B, and log-rank test with Bonferroni’s multiple comparisons for panel C. *p < 0.05, ****p < 0.0001 from control.

We found that earlier treatment with Cyrene resulted in the greatest increase in lifespan, but that maximal lifespan extension could be obtained when treatment is initiated during development or as young adults (**Figure 5B**). When Cyrene treatment was initiated at day 8 or day 12 of adulthood, it resulted in a small but significant decrease in lifespan. When Cyrene treatment was initiated at the L1 stage, the maximum lifespan extension was achieved when treatment continued until day 12 of adulthood (**Figure 5B**). Interestingly, lifelong Cyrene treatment initiated at the L1 stage resulted in a slightly shorter lifespan than treatment discontinued at day 12, raising the possibility that prolonged exposure may have subtle late-life costs rather than having uniformly beneficial effects.

Finally, we examined the extent to which a transient 2-day treatment with Cyrene beginning at young adulthood could extend lifespan. We found that this short exposure to Cyrene during early adulthood is sufficient to produce a significant, albeit slightly reduced, survival benefit (**Figure 5C**). These results indicate that Cyrene exerts its longevity effects during a specific window spanning from the L1 larval stage to day 5 of adulthood. The persistence of benefits after treatment withdrawal suggests that Cyrene triggers systemic adaptations, rather than acting as a chronic pharmacological buffer.

### Cyrene extends lifespan partially independently of DAF-16

DAF-16/FOXO is a conserved transcription factor required for the longevity effects of multiple interventions, including reduced insulin/IGF-1 signaling, germline ablation, mild impairment of mitochondrial function and certain forms of dietary restriction ^21–23^. To test the extent to which Cyrene requires DAF-16 to promote lifespan extension, we assessed its effects on the lifespan of *daf-16(mu86)* null mutants. We found that Cyrene significantly extended lifespan in *daf-16* mutants but to a lesser extent than in wild-type worms (**Figure 6A,B**). This indicates that the mechanism of lifespan extension in Cyrene-treated worms is at least partially independent of DAF-16. Because disruption of DAF-16 shortens lifespan, it is unclear whether the diminished extension in *daf-16* mutants reflects a reduced contribution of DAF-16 to Cyrene’s effects, or simply the lower baseline lifespan of the mutant background.

**Figure 6.**
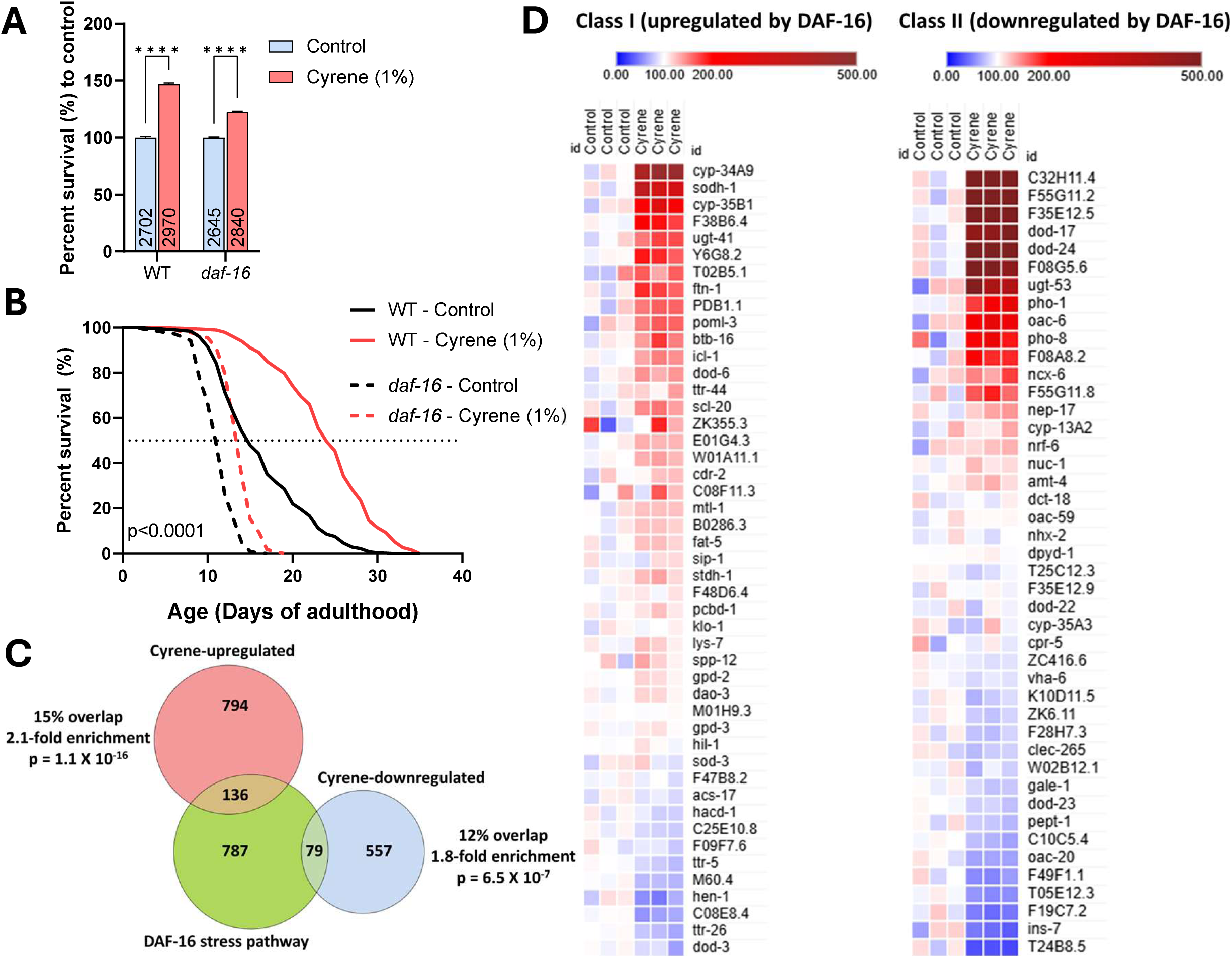
Cyrene extends lifespan in *daf-16* mutants and modulates DAF-16–associated transcriptional networks. (A) Normalized mean lifespans of *daf-16(mu86)* mutants treated with Cyrene from the L1 stage, demonstrating significant lifespan extension despite the absence of DAF-16, though the magnitude of effect is reduced compared to wild-type animals. (B) Corresponding survival curves showing the full lifespan distributions of Cyrene-treated and control *daf-16* mutants. (C) Overlap and enrichment analysis of genes differentially expressed in Day 8 adults treated with Cyrene from L1, showing that both upregulated and downregulated genes significantly intersect with DAF-16 stress-response targets, consistent with bidirectional modulation. (D) Heatmap displaying relative expression levels of top Class I (DAF-16–activated) and Class II (DAF-16–repressed) genes, indicating simultaneous up-and downregulation within DAF-16–regulated transcriptional modules. Error bars represent the standard error of the mean (SEM). Sample sizes (n-values) are indicated within each bar. Statistical analyses were performed using two-way ANOVA with Šidák’s multiple comparisons test in panel A, and log-rank test with Bonferroni’s correction in panel B. ****p < 0.0001 from control.

To explore this further, we qualitatively examined DAF-16::GFP localization in Cyrene-treated animals and observed no obvious differences in the nuclear localization of DAF-16 following treatment with Cyrene from L1 to early adulthood (data not shown). To determine the extent to which Cyrene treatment results in differential expression of DAF-16 target genes, we compared Cyrene-regulated genes (FDR < 0.05) from Day 8 adult worms to established target genes of the DAF-16 stress-response pathway ^16^. Interestingly, Cyrene treatment induced upregulated as well as downregulated genes, both were significantly enriched for DAF-16 targets (2.1-fold enrichment and 1.8-fold enrichment, respectively). This indicates that Cyrene has bidirectional effects on the expression of DAF-16 target genes (**Figure 6C**). This was further supported by examining the effect of Cyrene treatment on the expression of the top 50 DAF-16 upregulated (Class I) and downregulated (Class II) target genes ^24^. Cyrene treatment induced bidirectional regulation of both DAF-16-upregulated and DAF-16-downregulated genes (**Figure 6D**). Together, these results suggest that Cyrene does not strongly activate canonical DAF-16 signaling and that its ability to extend longevity is at least partially DAF-16 independent.

### Cyrene extends lifespan and modulates oxidative stress resistance in *Drosophila* in both sexes at multiple doses

To assess whether Cyrene’s pro-longevity effects are conserved across species, we tested its impact on lifespan in *Drosophila melanogaster*. Adult female and male flies were fed food containing Cyrene at defined concentrations ranging from 0.01% to 10% beginning in early adulthood (2 days post-eclosion) and their survival was quantified. Two independent trials were performed. In the first trial, we found that 10% Cyrene markedly decreases lifespan in both female and male flies and so this dose was not repeated in the second trial.

In females, lifespan extension was observed in both trials, with optimal effects at 0.1–1% (v/v) extending mean lifespan by 16–29% (**Figure 7A–C**). In males, the effect was concentration-specific, with 0.1% Cyrene consistently increasing mean lifespan significantly across both trials by 11–14%. While the survival curve reached statistical significance in one trial (p< 0.001), the other trial showed a strong trend toward extension (p=0.054) (**Figure 7D-F**).

**Figure 7.**
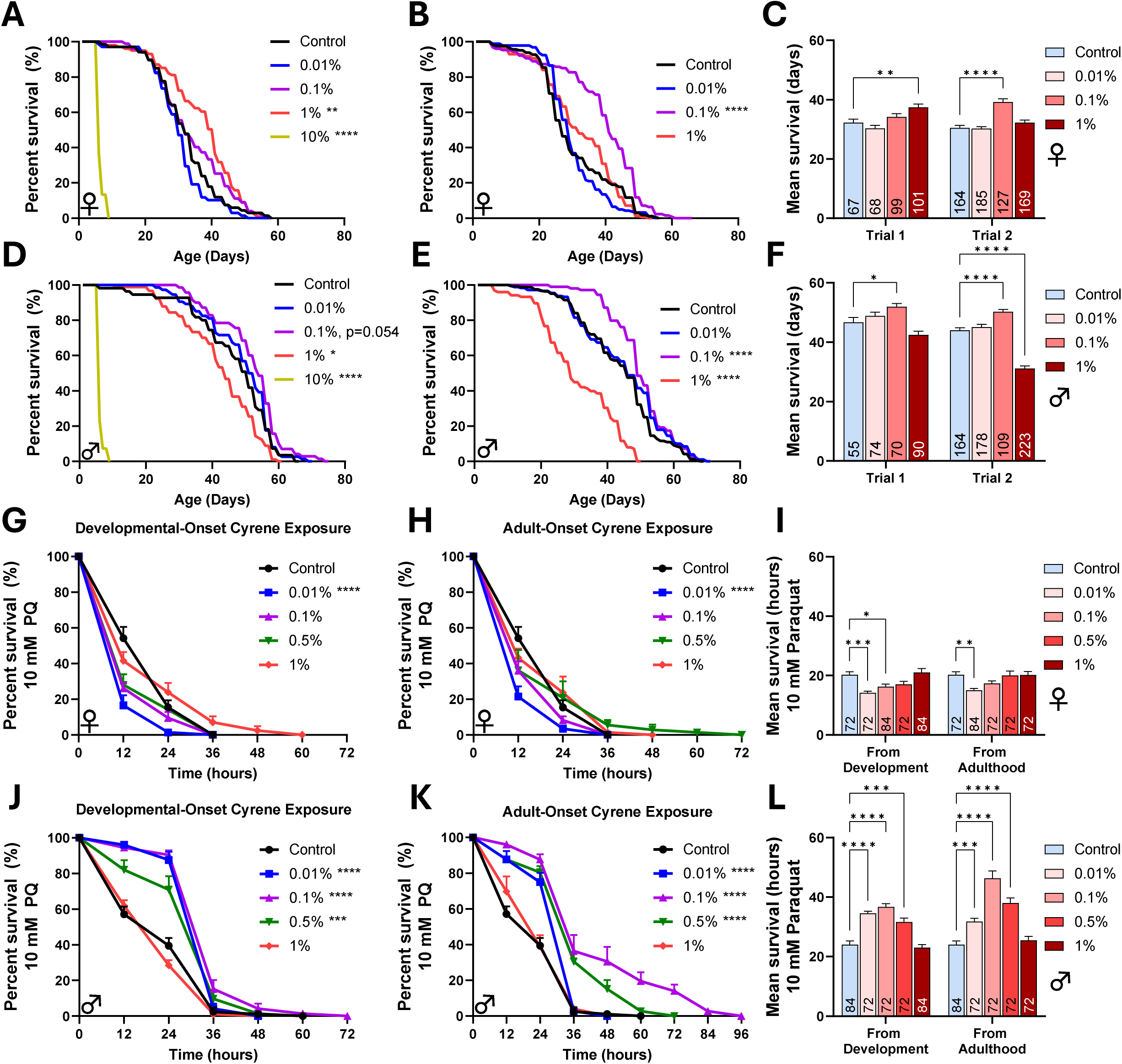
Cyrene extends lifespan and enhances stress resilience in *Drosophila* across sexes and at multiple doses. Panels (**A**) and (**B**) display survival curves from two independent trials in which adult female flies were exposed to increasing concentrations of Cyrene incorporated into their food, demonstrating dose-responsive lifespan extension. Panel (**C**) quantifies the corresponding mean lifespan values from each trial. Panels (**D**), (**E**), and (**F**) present the same experimental layout and analyses for male flies, revealing consistent benefits in both sexes. To assess the impact of Cyrene on oxidative stress resistance, Day 10 adult flies were exposed to 10 mM paraquat (PQ). In females, Cyrene was administered either (**G**) from the egg stage (developmental-onset) or (**H**) beginning in adulthood (adult-onset). Both conditions revealed reduced resistance to oxidative stress at lower Cyrene concentrations, with no significant difference at higher concentrations that extend lifespan under standard conditions. Panel (**I**) summarizes the corresponding mean survival data. Panels (**J**), (**K**), and (**L**) present the same experimental layout for male flies. In contrast to females, Cyrene exposure—whether initiated developmentally or in adulthood—enhanced resistance to paraquat in a bimodal, dose-dependent manner. Error bars represent the standard error of the mean (SEM). Sample sizes (n-values) are indicated within each bar. Statistical analyses were performed using the log-rank test with Bonferroni’s multiple comparisons correction in panels A, B, D, E, G, H, J and K, and two-way ANOVA with Dunnet’s multiple comparisons test in panels C, F, I and L. *p < 0.05, **p < 0.01, ***p < 0.001, ****p < 0.0001 from control.

Having shown that the ability of Cyrene to increase lifespan is conserved across species, we next examined the extent to which Cyrene could enhance resistance to stress in *Drosophila.* To do this, we treated flies with Cyrene beginning either at the start of development or at the start of adulthood and then challenged them with chronic oxidative stress using paraquat (10 mM) and assessed survival. In females, developmental or adult exposure to low Cyrene concentrations (0.01–0.1%) reduced resistance to paraquat, whereas higher concentrations (0.5–1%) had no effect (**Figure 7G–I**). In contrast, in males, both developmental-and adult-onset Cyrene exposure enhanced paraquat resistance in a bimodal, dose-dependent manner, with maximal protection observed at 0.1%, which nearly doubled mean survival under oxidative stress compared to control (**Figure 7J–L**).

Together, these findings demonstrate that Cyrene promotes lifespan extension and enhances biological resilience in both *C. elegans* and *Drosophila*, supporting the existence of conserved, cross-species mechanisms of action.

## Discussion

In this study we show that Cyrene is a novel, well-tolerated geroprotective compound that extends lifespan and healthspan in *C. elegans* and *Drosophila melanogaster*. We also show that Cyrene promotes resilience and protects against neurodegenerative disease, with minimal developmental or reproductive costs. Importantly, the beneficial effects of Cyrene are robust, reproducible, apply to both sexes and are conserved across species.

### Effect of chemical solvents on lifespan and physiological rates

Chemical solvents commonly used in *C. elegans* research can produce unintended physiological effects, depending on concentration, exposure window, and delivery method. DMSO (dimethyl sulfoxide), the most widely used solvent, has been reported to extend, reduce, or have no effect on lifespan—reflecting high context sensitivity across studies. In our work, we found that DMSO is well tolerated at concentrations ≤0.5%, with no measurable impact on lifespan, development, fertility, or locomotion under standard conditions ^25^. However, concentrations ≥0.5% acutely inhibit pharyngeal pumping and disrupt internal morphology, with effects shaped by genetic background and sensory processing capacity ^26^. At ≥0.75%, DMSO causes developmental delay and reduced brood size, while ≥2.5% markedly shortens lifespan. DMF (N,N-dimethylformamide) can extend lifespan by up to 50%, but its high toxicity severely limits its utility ^27^, and ethanol exhibits highly dose-and stage-specific effects ^28^.

In contrast, Cyrene extended lifespan by up to 75% in wild-type *C. elegans* without overt physiological disruption. A mild reduction in brood size was observed, but this was not accompanied by developmental delay or functional decline. Unlike conventional solvents, Cyrene not only avoids toxicity but also produces robust geroprotective effects, highlighting a unique biological activity that distinguishes it from typical vehicle controls.

### Cyrene as a novel lifespan-extending compound with healthspan benefits

Cyrene ranks among the most potent small-molecule geroprotectors identified in *C. elegans* to date. According to the DrugAge database ^29^, few compounds exceed a 50–60% increase in lifespan under standardized conditions. In our assays, Cyrene extended lifespan by up to 75% in wild-type animals, positioning it among the top of known longevity-promoting interventions (**Table 1**). Importantly, this magnitude of extension was achieved without the common trade-offs observed with many high-efficacy compounds, such as marked reductions in fertility or delayed development ^30,31^. This places Cyrene in a rare category of interventions that combine strong longevity effects with a favorable physiological profile.

**Table 1.** Lifespan-Extending Compounds in *C. elegans*: Comparative Magnitude of Effect. The table summarizes the maximal reported lifespan extension of the top 13 small-molecule geroprotective compounds from the DrugAge database. Cyrene is included for comparison, showing robust efficacy comparable to leading interventions.

| Compound | Concentration | Avg/Med Lifespan increase (%) | Reference |
| --- | --- | --- | --- |
| Cyrene | 1% v/v | Up to 75 | This study |
| Nitrophenyl piperazine containing compound 1 | 50 μM | 73 | Lucanic et al., 2016 <sup>55</sup> |
| Caffeine | 5 mM | 65 | Bridi et al., 2015 <sup>56</sup> |
| Rosmarinic acid | 180 μM | 63 | Lin et al., 2019 <sup>57</sup> |
| Resveratrol | 50 μM | 60 | Lim et al., 2018 <sup>58</sup> |
| Carbonylcyanide-3-chlorophenylhydrazone | 10 μM | 60 | Lemire, Behrendt, DeCorby, & Gaskova, 2009 <sup>59</sup> |
| Thioflavin T | 50-100 μM | 60 | Alavez, Vantipalli, Zucker, Klang, & Lithgow, 2011 <sup>60</sup> |
| Bacitracin | 0.01 | 59 | Lublin et al., 2011 <sup>61</sup> |
| Paraquat | 0.1 mM | 58 | Yang & Hekimi, 2010 <sup>62</sup> |
| Metformin | 25 mM | 57 | De Haes et al., 2014 <sup>63</sup> |
| Rifampicin | 30 μM | 56 | Golegaonkar et al., 2015 <sup>64</sup> |
| Blueberry extract | 60 μg/mL | 56 | Scerbak, Vayndorf, Hernandez, McGill, & Taylor, 2016 <sup>65</sup> |
| Salicylamine | 500 μM | 56 | Nguyen et al., 2016 <sup>66</sup> |
| Tanespimycin | 25 μM | 54 | Janssens et al., 2019 <sup>67</sup> |

The field of geroprotection has shifted from prioritizing lifespan extension alone to evaluating whether interventions can also enhance healthspan—the period of life spent in good functional condition. While foundational compounds such as rapamycin, resveratrol, and metformin have demonstrated the potential for pharmacological modulation of aging, each presents important translational considerations. Their efficacy often depends on variables such as dose, sex, species, and age of administration. Moreover, several interventions that extend lifespan in *C. elegans* do not consistently improve functional health and may, in some cases, extend the period of frailty.

For instance, canonical long-lived mutants like *daf-2*, *clk-1*, and *ife-2* increase lifespan but also prolong the proportion of life spent in a functionally compromised state ^8^. Axenic dietary restriction (ADR), while one of the most potent longevity interventions in worms, does not improve behavioral or functional metrics despite robust survival gains ^32^. Metformin, a compound of growing interest for translational geroscience, shows beneficial effects in young *C. elegans* but can be detrimental when administered later in life due to age-related loss of metabolic flexibility ^33^. These findings collectively highlight that longer lifespan does not inherently imply improved late-life function.

In this context, Cyrene stands out for its dual efficacy. It not only extends lifespan in *C. elegans* and *Drosophila* but also improves multiple healthspan parameters in *C. elegans*, including locomotor capacity, biological resilience and protection in models of tauopathy, amyloidosis, and polyQ aggregation. This balanced profile contrasts with several existing interventions where gains in longevity are accompanied by delayed growth, reduced fertility, or functional decline. Cyrene’s effects align with the goal of a proportionate extension of healthy life, rather than simply deferring decline.

This dual benefit supports the emerging consensus that lifespan extension alone is insufficient. Future geroprotectors must be evaluated for their ability to preserve robust physiological function across aging, including metrics such as mobility, stress response, and frailty onset ^34–38^. Our findings position Cyrene as a promising candidate that meets these evolving criteria, offering both longevity and functional preservation.

### Cyrene protects against neurodegenerative disease

Neurodegenerative diseases often involve converging cellular pathologies, including protein misfolding, oxidative stress, mitochondrial dysfunction, and impaired proteostasis ^39^. These features overlap significantly with the hallmarks of aging, suggesting that interventions promoting stress resilience may mitigate multiple forms of proteotoxicity ^40^.

Several longevity-promoting compounds have also shown benefit in neurodegenerative disease models ^2^. Metformin activates AMPK and reduces inflammatory and oxidative stress.

Resveratrol promotes mitochondrial function via SIRT1 activation. Rapamycin enhances autophagy and protein clearance through mTOR inhibition. Curcumin and N-acetylcysteine (NAC) are known for their antioxidant and anti-inflammatory effects. These compounds illustrate the diverse mechanisms through which longevity and neuroprotection can be achieved, including metabolic reprogramming, enhanced proteostasis, and redox balance.

Our results also show that Cyrene preserves locomotor function in *C. elegans* models of proteotoxic neurodegeneration. These improvements were seen across strains expressing human disease proteins including tau, α-synuclein, polyQ, and Aβ, suggesting that Cyrene acts not through disease-specific mechanisms, but by reinforcing shared cellular stress responses.

### Role of the microbial environment in mediating Cyrene’s effects

Drug–microbe interactions are a critical confounder in *C. elegans* aging studies. Several lifespan-extending compounds—including metformin—act indirectly by altering bacterial metabolism rather than targeting the host directly ^9,41^. Metformin, for example, promotes longevity by modulating microbial folate and methionine pathways, ultimately inducing a state of host methionine restriction, a conserved pro-longevity intervention across different species^42^.

By contrast, Cyrene’s effects appear largely independent of bacterial replication or metabolism. It does not impair bacterial growth at lifespan-extending concentrations and retains efficacy when worms are fed non-replicative or metabolically inactive bacteria, such as UV-killed or paraformaldehyde-fixed OP50. These findings suggest that Cyrene acts directly on the worm, bypassing common microbial confounds that limit the interpretability or translatability of many interventions ^13,14^. This distinction strengthens its candidacy as a bona fide geroprotective compound with host-intrinsic mechanisms of action.

### Mechanisms by which Cyrene promotes longevity

Lifespan assays with temporally restricted exposure revealed that early adulthood represents a critical window for Cyrene’s geroprotective effects. Short-term treatment during this period was sufficient to elicit long-lasting benefits, whereas late-life exposure produced diminished or even detrimental effects. These findings align with models of early-life hormesis, where transient stressors induce long-term physiological reprogramming that enhances late-life resilience ^43^. Notably, Cyrene conferred robust lifespan extension whether administered from the L1 larval stage or initiated at the onset of adulthood, prior to peak reproductive activity, indicating a broad window of efficacy.

This stands in contrast to several compounds identified in recent high-throughput screens, which extend lifespan only when administered during late larval development (L4) and lose efficacy when introduced in adulthood ^41^. For instance, LY-294002 and GSK2126458 failed to extend lifespan when treatment began at day 1 of adulthood (24 h after L4), despite being effective from L4. In contrast, Cyrene retained full potency when administered post-developmentally, suggesting its mechanism is not tightly coupled to developmental timing. This broader temporal flexibility distinguishes Cyrene from compounds with more restricted efficacy profiles.

DAF-16/FOXO is a central node in stress-responsive and longevity pathways, often required for lifespan extension triggered by reduced insulin/IGF-1 signaling, germline ablation, or metabolic stress ^21^. However, our findings demonstrate that Cyrene can extend lifespan even in the absence of DAF-16, indicating that its effects are not strictly dependent on this pathway.

However, the diminished response in *daf-16* mutants suggests the possibility of a partial contribution, rather than complete independence.

Although Cyrene did not trigger apparent DAF-16 nuclear localization during early adulthood, transcriptomic analysis in mid-life revealed enrichment of both activated and repressed DAF-16 target genes. This suggests that Cyrene modulates components of the DAF-16 transcriptional program later in life, potentially through indirect or compensatory mechanisms.

While the precise molecular mechanisms of Cyrene’s action remain to be defined, these findings support a model in which Cyrene reprograms aging trajectories by broadly enhancing organismal resilience. This distinguishes Cyrene from compounds such as myricetin and tyrosol, which require DAF-16/FOXO activity and fail to confer longevity benefits in its absence ^44,45^.

Notably, even DMSO has been shown to extend lifespan in a strictly DAF-16-dependent manner^46^, underscoring the unique DAF-16-independent properties of Cyrene.

Given the growing interest in combining pro-longevity interventions to maximize efficacy and circumvent pathway-specific limitations ^47^, future studies should explore how Cyrene interacts with other lifespan-extending compounds, particularly those acting independently of DAF-16 or through complementary stress-responsive mechanisms.

Identifying interventions that promote longevity through DAF-16–independent mechanisms, including mitochondrial signaling, chromatin remodeling, and proteostasis regulation, is critical for expanding the therapeutic landscape of aging. In this context, Cyrene emerges as a compelling candidate for further epistasis testing. Systematic analysis of its interactions with conserved stress regulators such as SKN-1, HSF-1, ATF-4, and components of the mitochondrial UPR may uncover additional dependencies or points of mechanistic convergence, further positioning Cyrene within the expanding class of stress-mimetic geroprotectors.

### Effects of Cyrene are conserved across species

Cyrene extends lifespan in both *C. elegans* and *Drosophila melanogaster*, demonstrating conserved efficacy across two phylogenetically distant invertebrates. In flies, lifespan extension was observed in both sexes, though optimal doses differed, potentially reflecting differences in uptake or metabolic context ^48–51^. Importantly, the route of exposure differs: while *C. elegans* experience continuous contact with Cyrene-supplemented agar, *Drosophila* are exposed only during feeding. These delivery differences likely influence internal dose and must be considered when comparing effects across species.

In addition to promoting longevity, Cyrene enhances oxidative stress resistance in both *C. elegans* and *Drosophila*, supporting conserved benefits across stress-response paradigms. In flies, this protective effect was more pronounced in males and most evident when exposure began in adulthood, consistent with our observations in *C. elegans* where early adulthood exposure was sufficient to trigger long-lasting benefits. This suggests that Cyrene acts during specific windows to induce adaptive reprogramming.

Together, these findings highlight Cyrene’s ability to promote longevity and stress resilience across species and contexts. Such cross-species conservation is a hallmark of robust geroprotective compounds. Curcumin, rapamycin, and resveratrol have shown similar benefits in nematodes, flies, and in some cases, mammals ^2^. Cyrene’s consistent effects across evolutionary and phenotypic boundaries support its engagement of core aging-related pathways and reinforce its candidacy for mechanistic dissection and evaluation in vertebrate models.

## Conclusion

In this work, we show that Cyrene is a novel geroprotective compound that extends lifespan and healthspan in *C. elegans* and *Drosophila melanogaster*. Cyrene enhances stress resilience in wild-type animals and preserves locomotor function in models of neurodegeneration. The effects of Cyrene are dose-dependent, require early-life exposure, are independent of bacterial metabolism and are at least partially independent of *daf-16*. Overall, this work demonstrates that Cyrene is a conserved, well-tolerated modulator of aging and highlights the potential of non-canonical small molecules to reprogram resilience and longevity across species.

## Methods

### Strains

The strains were maintained on nematode growth medium (NGM) plates at 20°C unless otherwise stated, with OP50 *Escherichia coli* bacteria as the nutritional source. The following *C. elegans* strains were obtained from the *Caenorhabditis* Genetics Center (CGC): N2 (wild-type), CF1038 *daf-16(mu86)*, CL2355 *dvIs50* [*snb-1p::Aβ₁₋₄₂*; *mtl-2p::GFP*], GRU102 *gnaIs2* [*myo-2p::YFP*; *unc-119p::Aβ₁₋₄₂*], JVR182 (*dat-1p::α-syn(A53T)*; *cwrIs856 [dat-1p::GFP; dat-1p::LRRK2(G2019S); lin-15(+)*]; *ges-1p::RFP)*, JVR389 [*eft-3p::α-syn::RFP*; *dat-1p::GFP*], NL5901 *pkIs2386* [*unc-54p::α-syn::YFP*; *unc-119(+)*]. AM716 *rmIs284* [*rgef-1p::Q67::YFP*] was kindly provided by R. I. Morimoto, CK10 *bkIs10 [aex-3p::tau(h4R1N V337M); myo-2p::GFP]* was kindly provided by B. C. Kraemer, MQ1698 [*unc-54p::Htt74Q::GFP*] was kindly provided by M. J. Monteiro.

### Bacterial Culture Preparation

*Escherichia coli* strain OP50 was used as the standard food source. Cultures were initiated from glycerol stocks streaked onto LB agar plates and incubated overnight at 37 °C. A single colony was grown in 250 mL of LB broth in a 1 L Erlenmeyer flask at 37 °C for 20 hours with shaking.

Cultures were centrifuged at 3900 x *g* for 10 minutes at room temperature, and the bacterial pellet was resuspended in a reduced volume of supernatant to generate concentrated stocks.

### Synchronization of Worms

Gravid adults were cultured on 6 cm or 10 cm NGM plates seeded with 2X concentrated OP50 *E. coli* (600 µL and 1 mL per plate, respectively) until sufficient egg-laying occurred. Worms and laid eggs were collected in M9 buffer (22 mM KH₂PO₄, 42 mM Na₂HPO₄, 86 mM NaCl, 10 mM MgSO₄), transferred to 15 mL tubes, and centrifuged at 1300 × g for 2 min. After one M9 wash, 250 µL of supernatant was left above the pellet, and 2 mL of freshly prepared bleach solution (1% sodium hypochlorite, 625 mM NaOH) was added. Samples were vortexed for 1 min and rested for 30 s, repeated as needed (up to 6 min total) until adult carcasses were dissolved. The suspension was washed three times with M9 and resuspended in 2 mL fresh M9. Eggs were incubated at 20°C on a rotating platform for ∼20 hours to allow synchronous hatching in the absence of food. Synchronized L1 larvae were quantified by microscopy and transferred to fresh NGM plates for experiments.

### Cyrene Plate Preparation

Standard NGM was prepared and supplemented with 50 µM FUdR when required. Plates were poured with 2.5 mL agar per 3.5 cm plate or 6 mL per 6 cm plate, dried overnight at room temperature, and seeded with 2X OP50 (250 µL for 3.5 cm, 600 µL for 6 cm). Seeded plates were dried for 3 days and stored inverted at 4°C. For treatment, Cyrene (Sigma-Aldrich, 807796) or vehicle (ddH₂O) was applied the day before use. Fresh 10X stock solutions were added on top of the agar (250 µL for 3.5 cm, 600 µL for 6 cm plates), mixed on a nutator (70 RPM, 5–10 min), and incubated in an air-tight plastic box overnight at 20°C to equilibrate. Plates with incomplete absorption or lawn disruption were excluded. An in-agar method, in which Cyrene was added directly to molten NGM prior to pouring, was also tested. Although both methods extended lifespan, the on-top agar approach yielded stronger and more consistent effects and was used for all subsequent experiments.

### Brood size

Brood size was determined by placing individual L4 worms on NGM plates. Worms were transferred daily to new plates until egg laying ceased. The progeny was allowed to develop up to at least the L3 stage before quantification. Three biological replicates were performed for each strain, with each biological replicate containing at least four worms.

### Post-embryonic Development

Postembryonic development (PED) was assessed by transferring synchronized arrested L1s to agar plates. Starting at 24 hours after transferring, worms were scored for their developmental stage. Four biological replicates of 25 animals minimum each were completed.

### Thrashing rate

Adult worms cultured on 60 mm NGM plates seeded with OP50 *E. coli*, and a volume of 2 mL of M9 buffer was added to the worms. The worms were left to acclimatize for 1 min before capturing 1 min video at 14 FPS using WormLab (MBF Bioscience). The videos were analyzed using wrMTrck plugin for the open-source image processing software, Fiji. Worm tracks were only considered for analysis when they included at least 420 frames (30 seconds). At least three biological replicates were completed for each condition.

### Lifespan assay

All lifespan assays were performed at 20°C. Lifespan assays included 50 µM 5-fluoro-2’-deoxyuridine (FUdR) to limit the development of progeny and the occurrence of internal hatching. This concentration of FUdR completely prevents the development of progeny to adulthood and does not affect the lifespan of wild-type worms ^52^. Animals were excluded from the experiment if they crawled off the plate, burrowed or displayed internal hatching (matricide) or vulval rupture. However, these censored worms were still included in subsequent statistical analysis for lifespan. At least four biological replicates were completed for each condition.

### Preparation of Non-Replicative Bacterial Lawns

To assess whether Cyrene’s effects depend on bacterial viability, *E. coli* OP50 was rendered non-viable by either UV-C irradiation or paraformaldehyde fixation.

### UV-killed bacteria

NGM plates seeded with *E. coli* OP50 were exposed to UV-C light (254 nm, 6.4 J/cm² over 30 minutes) using a UV transilluminator and stored at 4 °C. Immediately before Cyrene treatment, plates received an additional 15-minute UV-C exposure (3.2 J/cm²) to ensure complete bacterial inactivation. Five biological replicates were completed for each condition.

### Paraformaldehyde-fixed bacteria

Metabolically inactive bacteria were prepared as previously described with a modification ^14^. Briefly, OP50 cultures were grown overnight in LB, harvested by centrifugation, and washed with PBS. Bacterial pellets were resuspended in PBS containing 0.5% methanol-free formaldehyde and incubated at 37°C for 1 hour with shaking. Mock-treated controls were processed identically without fixative. After fixation, bacteria were washed thoroughly and adjusted to optical density (OD₆₀₀ ∼ 6.5). NGM plates (6 cm, ±50 µM FUdR) were seeded with 600 µL of the suspension and dried before use in lifespan assays.

### Heat stress resistance

NGM plates (6 cm) were seeded with 75 µL of 10X concentrated OP50 bacteria. Seeded plates were allowed to dry at 35°C for ∼15 minutes. Once dried, adult worms were transferred to the plates and incubated at 35°C for 3 or 6 hours, after which plates with worms were transferred to 20°C. Scoring for survival occurred 24 hours after the start of the heat shock (24 hours after the worms were placed into the 35°C). Five biological replicates were completed for each condition.

### Osmotic stress resistance

NGM plates containing 450 mM or 500 mM NaCl were poured and allowed to dry on the bench before storing at 4°C. One day before the experiment, 200 µL of 5X concentrated OP50 *E. coli* was seeded onto each plate and allowed to dry overnight. The next day, adult worms were picked into the hypertonic NGM plates and incubated at 20°C for 48 h before scoring survival. Worms with internal hatching were censored and excluded from the total number of deaths. At least four biological replicates were completed for each condition.

### Resistance to Acute Oxidative Stress

Acute oxidative stress resistance was assessed using juglone and paraquat exposure assays. **Juglone assay:** Fresh juglone (dissolved in ethanol) was added to autoclaved, cooled NGM to final concentrations of 140–440 µM. The medium was poured into 60 mm plates, seeded with *E. coli* OP50, and dried briefly before use. Adult worms were transferred to plates and scored for survival every 2 hours for up to 6 hours. Four biological replicates were completed for each condition.

### Paraquat assay

For acute paraquat stress, NGM containing 100 µM FUdR was supplemented with 220 mM paraquat (Methyl viologen dichloride hydrate). Plates were poured, dried, and seeded with OP50 the day before use. Adult worms were exposed the following day, and plates were used within 48 hours of preparation to ensure consistency. Four biological replicates were completed for each condition.

### Resistance to chronic oxidative stress

Resistance to chronic oxidative stress was assessed by exposing worms to 4 mM or 10 mM paraquat. Paraquat freshly dissolved in ddH_2_O was added to NGM before pouring and 100 µM FUdR was included in the plates to prevent internal hatching that is caused by exposure to paraquat. The plates were allowed to dry overnight before storage at 4°C. One day before the experiment, the paraquat plates were seeded with 200 µL of 10X concentrated OP50 *E. coli* and allowed to dry overnight. The next day, adult worms were transferred to these plates and survival was monitored every day until death. Four biological replicates were completed for each condition.

### UV genotoxic stress assay

Age-synchronized worms at defined adult stages were transferred to NGM plates containing 50 µM FUdR and seeded with *E. coli* OP50. Plates were exposed to 0.1 J/cm² of UV-C light (254 nm) using a UV transilluminator. After irradiation, plates were returned to 20 °C, and worms were monitored daily for survival. Death was scored based on the absence of response to gentle mechanical stimulation. Four biological replicates were completed for each condition.

### Dithiothreitol (DTT) reductive stress assay

Fresh DTT stock (500 mM in ddH₂O) was prepared on the day of use and protected from light. DTT was added to molten NGM (containing 50 µM FUdR) to final concentrations of 1, 2.5, or 5 mM. Plates were poured, seeded with *E. coli* OP50, and dried before use. Worms were transferred at Day 1 or Day 8 of adulthood and scored daily for survival. Four biological replicates were completed for each condition.

### Tunicamycin endoplasmic reticulum stress assay

Tunicamycin stock (10 mg/mL in DMSO) was freshly prepared and added to molten NGM (with 50 µM FUdR) to achieve final concentrations of 25 or 50 µg/mL. Plates were poured, protected from light, and stored at 4°C. Prior to use, plates were seeded with OP50 and dried overnight. Worms were transferred at Day 1 or Day 8 of adulthood and monitored daily for survival. Four biological replicates were completed for each condition.

### Bacterial growth inhibition assay

To assess the bactericidal potential of Cyrene, a dose-response assay was performed using *E. coli* OP50. Overnight cultures grown in LB medium at 37°C were diluted to a starting OD₆₀₀ of 0.1 in fresh LB. Cultures were aliquoted in and treated with various concentrations of Cyrene or DMSO (vehicle control). Water was used as an untreated control, and 100 µg/mL carbenicillin served as a positive control. Tubes were incubated at 37°C for 2 hours with shaking at 250 RPM. OD₆₀₀ values were measured in triplicates to assess bacterial growth. Three biological replicates were completed for each condition.

### RNA extraction

Synchronized *C. elegans* (∼1500 per replicate) were harvested from NGM plates into 15 mL centrifuge tubes using M9 buffer. Worms were allowed to settle by gravity for 3 minutes or by centrifugation at 100 × g for 1 minute, then washed three times with M9 to remove residual bacteria. After the final wash, ∼200–300 µL of buffer was left above the worm pellet, and 1 mL of cold TRIzol reagent (Thermo Fisher) was added directly without disturbing the pellet.

Samples were snap-frozen in liquid nitrogen and stored at −80 °C until lysis.

For homogenization, TRIzol-suspended worms were transferred to pre-chilled Zymo BashingBead Lysis Tubes (0.1/0.5 mm beads; Zymo Research) and vortexed at 3000 RPM using a horizontal tube holder (Fisher Scientific). Lysis was performed for up to 4 minutes with intermittent cooling (e.g., 1–2 minutes vortexing followed by 1 minute on ice). Samples were centrifuged at 12,000 × g for 1 minute, and the supernatant containing total RNA was transferred to RNase-free microcentrifuge tubes and stored at −80 °C.

Total RNA was purified using the Direct-zol RNA Miniprep Kit (Zymo Research) according to the manufacturer’s instructions.

### RNA sequencing analysis

RNA sequencing was performed by Genome Quebec. Each RNA-seq sample consisted of pooled RNA from two independent biological replicates mixed at equal concentrations. This pooling strategy was applied consistently across all experimental conditions to reduce individual variability while preserving biological relevance ^53^. All samples were processed in parallel to minimize batch effects during library preparation and sequencing.

Total RNA underwent standard quality control assessments, including integrity and concentration measurements. Poly(A)-enriched libraries were constructed and sequenced on an Illumina NovaSeq 6000 platform using paired-end 100 bp reads (PE100), generating ∼25 million reads per sample.

Bioinformatic analysis was performed by the McGill Genome Centre. Adaptor trimming and low-quality base removal (Phred < 30) were performed using Trimmomatic (Bolger et al., 2014). Reads were aligned to the *C. elegans* reference genome (WBcel235) using STAR (Dobin et al., 2012), and gene-level counts were obtained with HTSeq (Anders et al., 2015) using the parameters-m intersection-nonempty-stranded=reverse. Genes with an average read count <10 across all samples were excluded. Raw counts were normalized using edgeR’s TMM method (Robinson et al., 2010) and transformed into log2 counts per million (log2CPM) using the voom function from limma (Ritchie et al., 2015). Differential expression analysis was performed using a linear model fitted with lmFit, and *p*-values were adjusted for multiple testing using the Benjamini–Hochberg procedure. Gene set enrichment analysis was carried out using fgsea (http://bioconductor.org/packages/fgsea/) on genes ranked by *t*-statistic.

### Lifespan validation in *Drosophila*

Wild-type *Drosophila melanogaster* (Canton-S) were reared at 25 °C on standard cornmeal-based fly food. Progeny were collected upon eclosion and allowed to mate under standard conditions for 48 hours. Males and females were then separated, and 25–27 flies of a given sex were transferred into individual glass vials containing food supplemented with either Cyrene or vehicle control (water). Cyrene treatment therefore began two days post-eclosion, following the completion of mating. Flies were transferred to fresh food every other day, and survival was recorded at each transfer until all individuals died. Flies that escaped, became trapped in the food, or died due to handling were censored from the analysis. Each condition was tested using 7–10 biological replicates per sex.

### Oxidative stress resistance assay in *Drosophila*

To assess oxidative stress resistance, two groups of *Drosophila melanogaster* (Canton-S) were established. In the first group (developmental-onset), adult flies were allowed to mate and lay eggs on food supplemented with varying concentrations of Cyrene. Offspring from this group were exposed to Cyrene continuously throughout development and adulthood. In the second group (adult-onset), mating and egg laying occurred on standard food, and progeny were transferred to Cyrene-supplemented food only after development was complete.

Adult progeny from both groups were collected on the same day and aged for 10 days on their respective Cyrene-supplemented diets. On Day 10, flies were transferred to oxidative stress assay food containing 1% agar, 5% sucrose, and 10 mM paraquat (PQ). For each condition, 6–7 replicate vials containing 12 flies each were tested. Mortality was recorded twice daily until all flies died.

### Statistical analysis

Statistical analysis was performed using GraphPad Prism version 9, and log-rank (Mantel-Cox) tests were performed using online application for survival analysis 2 (OASIS 2)^54^. Statistical tests utilized are indicated in the figure legends. Error bars indicate standard error of the mean (SEM). ns=not significant, *p<0.05, **p<0.01, ***p<0.001, ****p<0.0001.

## Acknowledgments

Some strains were provided by the CGC, which is funded by NIH Office of Research Infrastructure Programs (P40 OD010440). We would also like to acknowledge the *C. elegans* knockout consortium and the National Bioresource Project of Japan for providing strains used in this research. This work was supported by the Canadian Institutes of Health Research (CIHR; http://www.cihr-irsc.gc.ca/; JVR), the Natural Sciences and Engineering Research Council of Canada (NSERC; https://www.nserc-crsng.gc.ca/index_eng.asp; JVR), the National Institute on Aging (NIA, R01 AG082801, R01 AG082696; AP), and the National Institute of General Medical Science (NIGMS, R35 GM146869; AP). JVR is the recipient of a Senior Research Scholar career award from the Fonds de Recherche du Québec Santé (FRQS) and Parkinson Quebec. The funders had no role in study design, data collection and analysis, decision to publish, or preparation of the manuscript. We thank Jatin Malhotra for assistance with *Drosophila* lifespan assays and media preparation.

## Author Contributions

Conceptualization: AA, JVR. Methodology: AA, SY, AP, AAP, JVR. Investigation: AA, SY, AP. Analysis: AA, SY, AP, AAP, JVR. Validation: AA, SY, AP, AAP, JVR. Visualization: AA, SY, AP, AAP, JVR. Writing – original draft: AA. Writing – review and editing: AA, SY, AP, AAP, JVR. Supervision: JVR, AAP.

## Data and materials availability

Statistical analyses for post-embryonic development are provided in Supplemental Table S1. Raw lifespan data are available in Supplemental Table S2. Other raw data will be provided upon request. All materials used in this manuscript are available to be shared with the scientific community. Requests for data or materials should be addressed to Jeremy Van Raamsdonk.

## Supplementary Figures

**Supplementary Figure 1.**
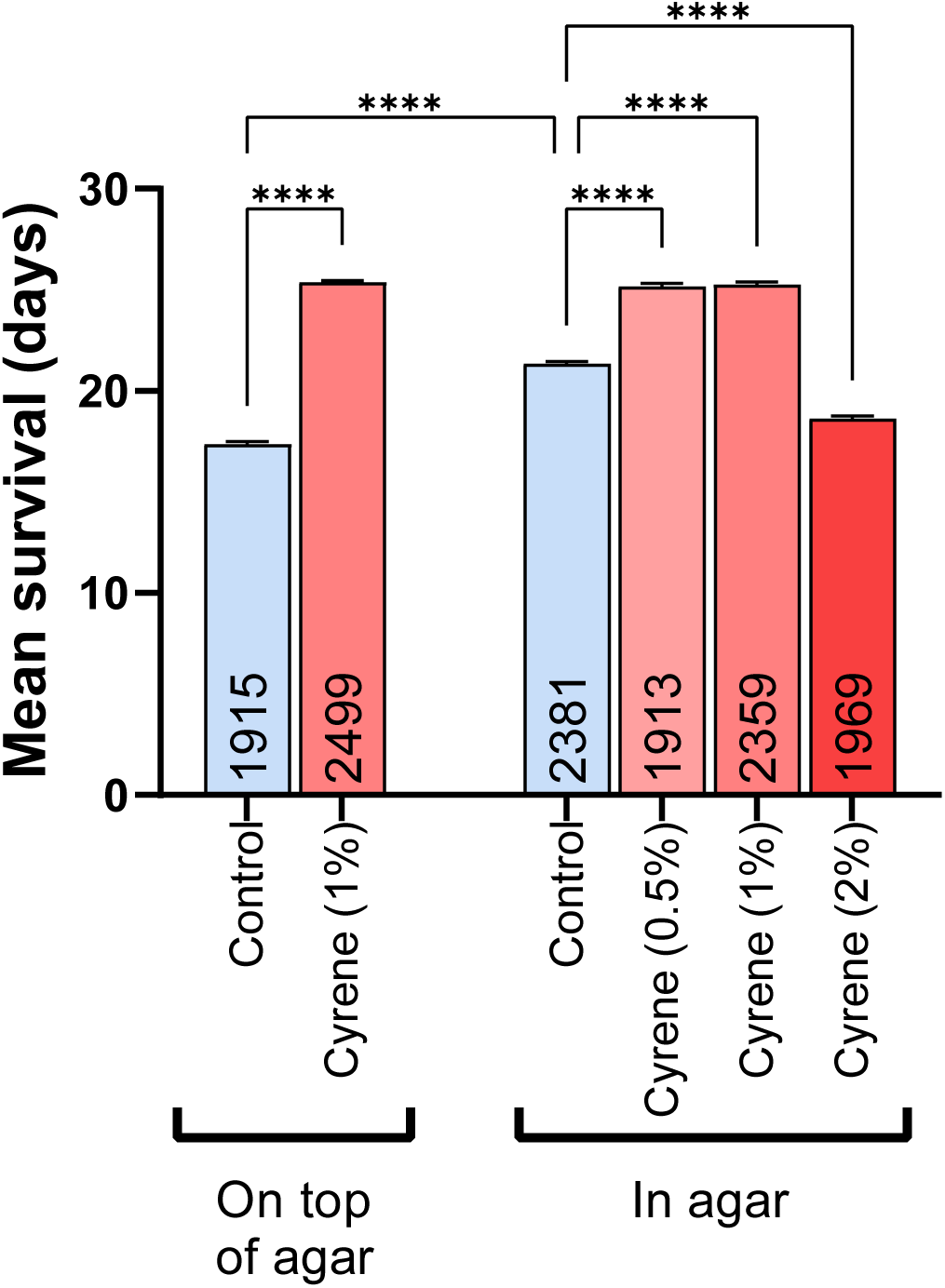
Cyrene increases lifespan independent of method of administration. Lifespan of *C. elegans* treated with Cyrene applied either on-agar (post-seeding) or in-agar (pre-seeding, during plate preparation), showing a consistent longevity benefit across both delivery methods. Error bars indicate SEM. Sample sizes (n-values) are indicated within each bar. Statistical analysis was performed using two-way ANOVA with Tukey’s multiple comparisons test. ****p<0.0001.

**Supplementary Figure 2.**
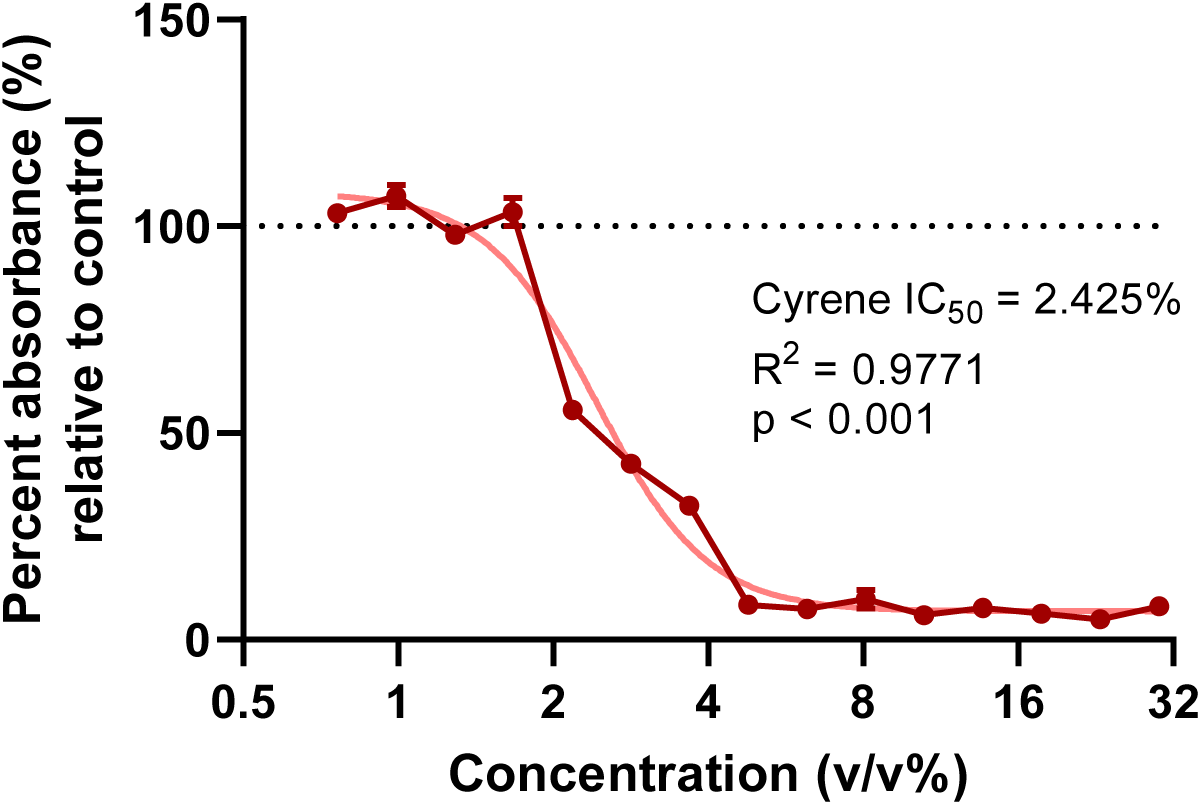
Cyrene inhibits bacterial growth at concentrations above 2%. Dose–response curves of *E. coli* OP50 growth following exposure to Cyrene or DMSO. Bacterial density (OD₆₀₀) was measured and fitted using non-linear regression. The half-maximal inhibitory concentration (IC₅₀) and coefficient of determination (R²) are indicated. Error bars represent the standard error of the mean (SEM).

**Supplementary Figure 3.**
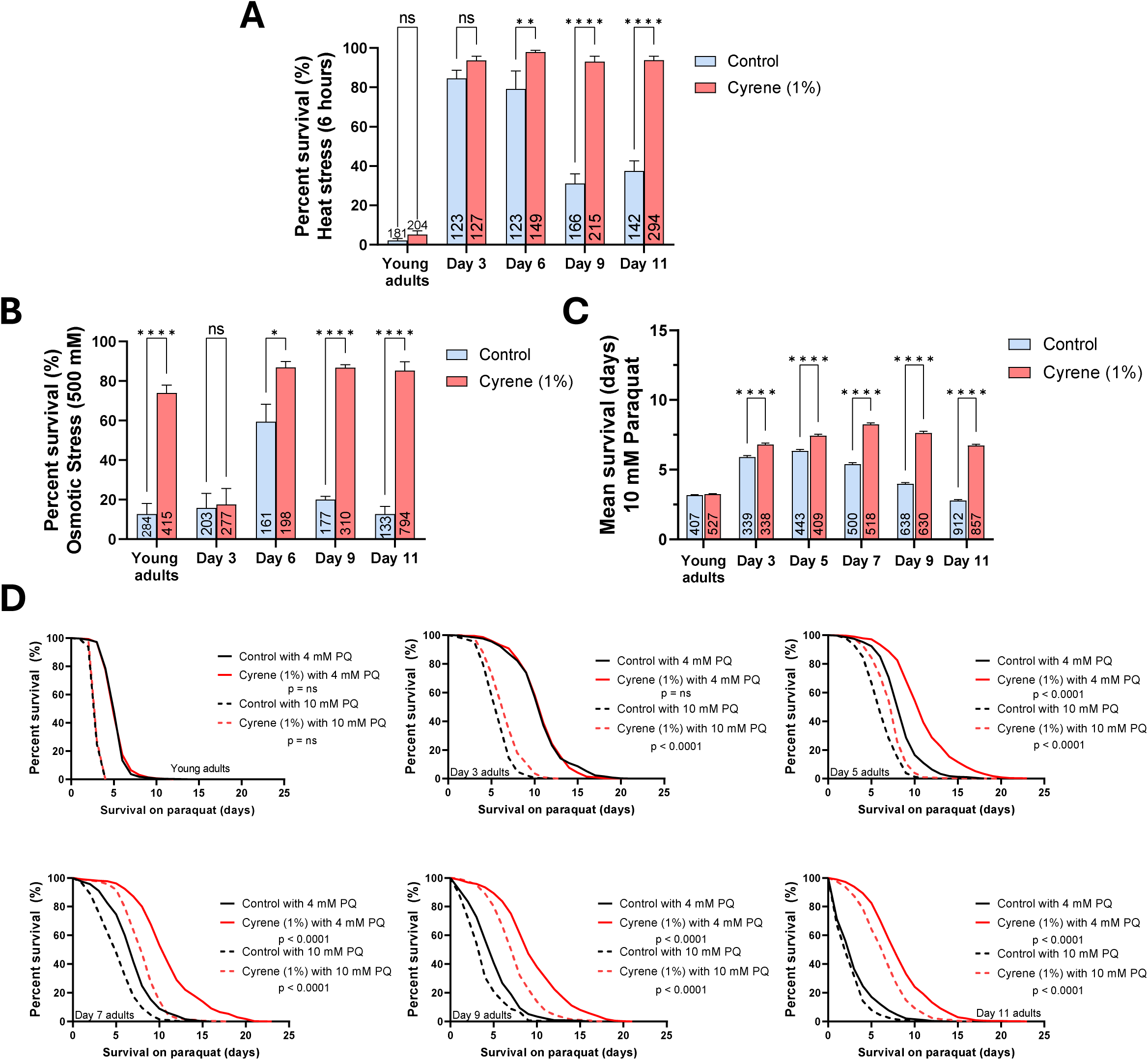
Cyrene promotes survival under diverse acute and chronic stress conditions. Cyrene-treated *C. elegans* exhibited enhanced resilience across multiple environmental stressors, suggesting broad activation of cellular stress defense pathways. (A) Under sub-lethal heat stress (35°C for 6 hours followed by 18 hours of recovery), Cyrene preserved survival capacity in mid-to-late adulthood, while stress resistance in control animals declined with age. (B) In hypertonic conditions (500 mM NaCl for 48 hours), Cyrene conferred significantly improved survival relative to controls. (C) Cyrene treatment also improved survival under chronic oxidative stress induced by 10 mM paraquat. (D) Survival curves across two paraquat concentrations (4 and 10 mM) confirmed the protective effect, with Cyrene-treated animals consistently exhibiting higher stress resistance. Error bars represent the standard error of the mean (SEM). Sample sizes (n-values) are indicated within each bar. Statistical analyses were performed using two-way ANOVA with Šidák’s multiple comparisons test in panels A-C, and log-rank test with Bonferroni’s multiple comparisons in panel D. ns = not significant, *p < 0.05, **p < 0.01, ****p < 0.0001.

**Supplementary Figure 4.**
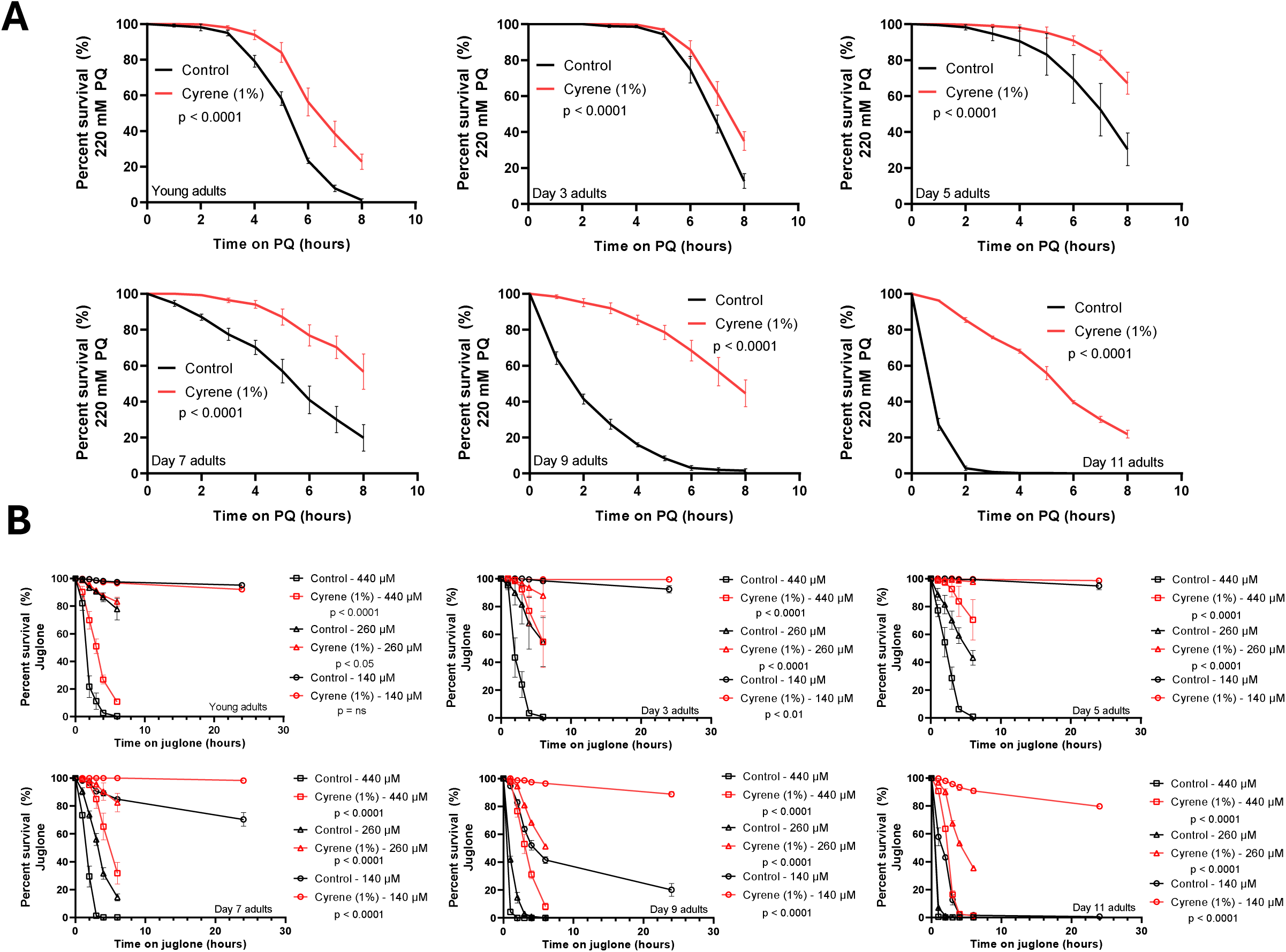
Cyrene increases resistance to acute oxidative stress. Cyrene-treated *C. elegans* exhibited enhanced survival in response to two distinct acute oxidative stressors. (A) Exposure to high-dose paraquat (220 mM) revealed a significant survival advantage in Cyrene-treated animals compared to controls. (B) Cyrene also improved survival under acute juglone-induced oxidative stress across multiple concentrations (140, 260, and 440 µM). Error bars represent the standard error of the mean (SEM). Statistical analyses were performed using the log-rank test with Bonferroni’s multiple comparisons.

**Supplementary Figure 5.**
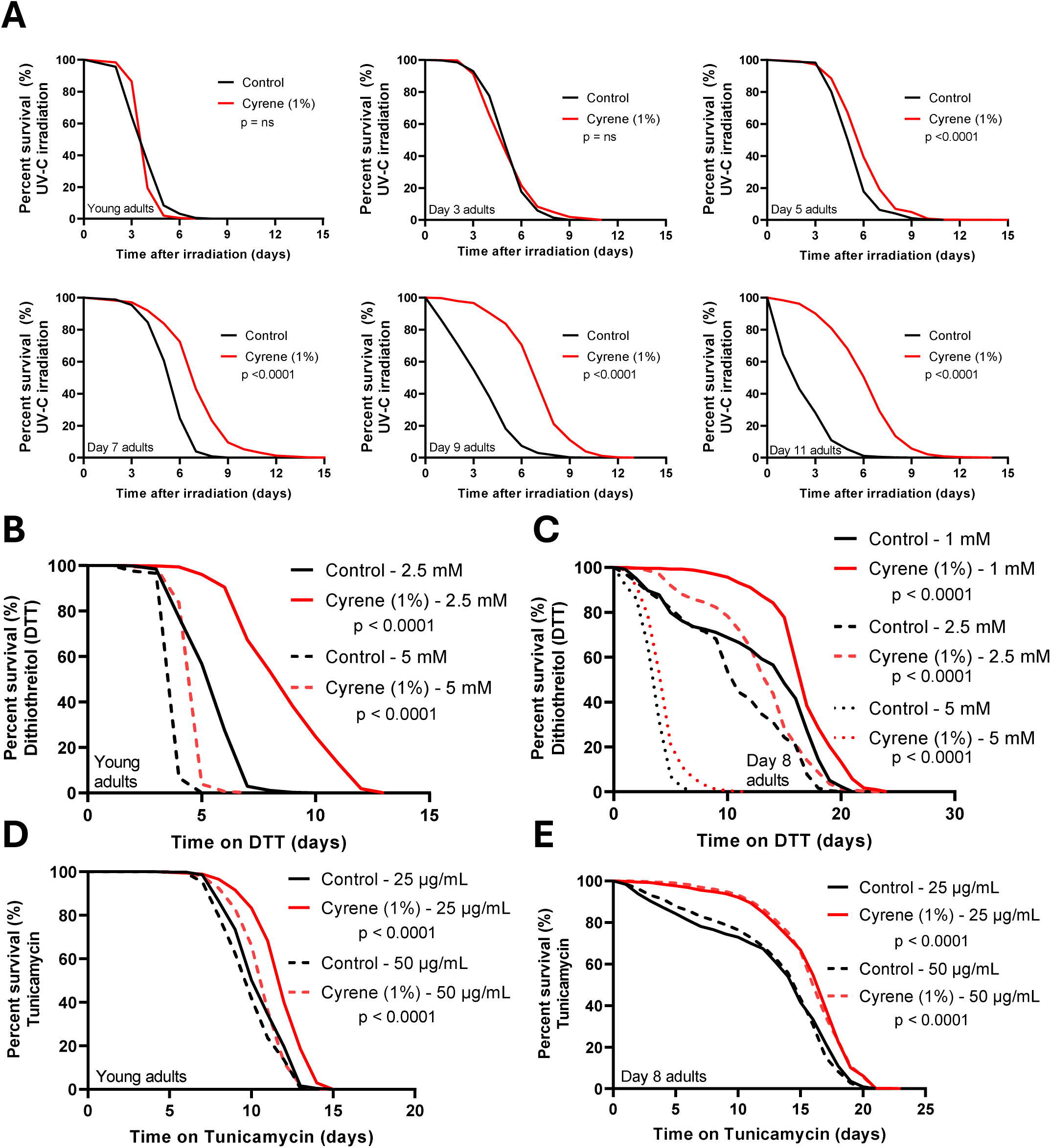
Cyrene enhances resistance to genotoxic, thiol-reductive, and ER stress. Cyrene-treated *C. elegans* exhibited improved survival under diverse acute stress conditions beyond oxidative damage. (**A**) Cyrene conferred increased survival following UV-C irradiation (0.1 J/cm²), indicating enhanced resistance to genotoxic stress. (**B–C**) Resistance to thiol-reductive stress induced by dithiothreitol (DTT) was significantly improved in both young (**B**) and day 8 adult (**C**) animals. (**D–E**) Similarly, Cyrene increased survival under tunicamycin-induced ER stress in both young (**D**) and aged (**E**) worms, suggesting preserved proteostasis with age. Young adult and day-8 time points were selected to capture early and later stages of adulthood, reflecting potential age-associated shifts in stress response capacity observed in preliminary assays. Statistical analyses were performed using the log-rank test with Bonferroni’s multiple comparisons.

